# Synthetic pectin-cellulose nanofiber capsule provides minimal model capturing mechanics of a regenerating plant cell wall

**DOI:** 10.1101/2025.10.07.680914

**Authors:** Cyril Grandjean, Ravi Shanker, Sarah A. Pfaff, Anran Mao, Jordi Chan, Sophie Asnacios, Atef Asnacios, Sulin Zhang, Daniel J. Cosgrove, Enrico Coen, Anna J. Svagan, Pauline Durand-Smet

**Affiliations:** Université Paris Cité, CNRS UMR 7057, Laboratoire Matière et Systèmes Complexes, Paris, France; Royal Institute of Technology, KTH, Teknikringen 56-58, SE-100 44, Stockholm, Sweden; Pennsylvania State University, Department of Biology, University Park, Pennsylvania 16802, United States; John Innes Centre, Department of Cell and Developmental Biology, Norwich Research Park, Colney Lane, Norwich NR4 7UH, UK

**Keywords:** Mechanical properties, primary cell wall, protoplasts, cellulose microfibrils, layer-by-layer, synthetic analogues, cellulose nanofibers

## Abstract

Plant primary cell walls are dynamic supramolecular assemblies composed of layered cellulose, hemicellulose and pectin, progressively built through synthesis and secretion. However, the specific architectural features and structural components sufficient to endow the mechanical properties of the wall remain unclear. Here, we construct a minimal synthetic spherical shell and compare its structural and mechanical properties to those of a plant single-cell system. To eliminate complexities from intercellular connectivity and developmental history, we exploit the ability of plant protoplasts to regenerate cell walls de novo. Compression tests of regenerating protoplasts between parallel plates reveal that wall stiffness increases with wall thickening over time. Despite differences in assembly pathways, architecture, and composition, the synthetic shell exhibits a similar thickness-dependent modulus and similar material stiffness. The synthetic shell, composed of pectin and cellulose layers, mirrors the mechanical behavior of regenerating primary cell walls, suggesting that these components are sufficient to confer key viscoelastic properties in the limit of small deformations. Given the complexity of natural plant cell walls, the synthetic analogue offers a controllable platform to dissect the mechanical contributions of individual wall components.

## 1. Introduction

Plant primary cell walls are composed of multiple layers arranged in a cross-lamellate architecture. Each layer consists of interwoven polysaccharide networks of cellulose, hemicellulose and pectin that are progressively assembled through cellulose synthesis at the plasma membrane and matrix secretion from Golgi-derived vesicles (1–7). Because the wall deposition occurs sequentially, its composition and architecture are shaped not only by developmental history (8) but also by mechanical stresses imposed during growth (9–14). The result is a heterogeneous complex three-dimensional hydrated polymer network (15).

This structural complexity raises a fundamental question: which features of the wall contribute to its mechanical resilience versus other aspects such as growth, signaling, metabolite transport, or protection from pathogens and environmental stressors (16, 17). One approach to address this question is through genetic or enzymatic perturbations, using mutants defective in wall biosynthesis or applying enzymes that degrade or modify specific wall polymers (18–23). These studies have provided valuable insight into the role of various wall components (pectin, cellulose and xyloglucan), by quantifying how these perturbations modify aspects of wall mechanics such as elasticity, viscoelasticity, loosening or softening under various loading conditions. However, while such studies can identify necessary components that contribute to wall mechanics, they do not establish sufficiency. A synthetic approach provides a complementary strategy: one aimed at identifying the minimal set of components sufficient to recapitulate specific mechanical properties. Here, we focused on the wall’s viscoelasticity and its link with wall thickness and layering.

Early efforts to synthetically mimic wall mechanics involved cellulose-based bacterial films made of bacterial cellulose nanoribbons combined with hemicellulose or pectin (24, 25). However, these composites showed much lower stiffness (26, 27) than native plant cell walls (28), likely due to missing structural features such as fiber alignment, lamellation, nanoscale thickness (≈ 30 - 50 nm for native walls vs. ≈ 2-3 nm for synthetic composites), and proper cellulose-matrix interactions (29, 30). An alternative strategy involves layer-by-layer assembly of capsules made of alternating nanolayers of wood-derived cellulose nanofibers and pectin, a method that offers precise control of composition and thickness (31). Although such capsules have previously been reported (32–34), their mechanical properties remain uncharacterized, and no direct comparison has been made with native cell walls.

A convenient method for assessing the mechanical properties of synthetic capsules is parallel-plate compression (35), a technique applied to both animal and plant cells (36–38). Here, we use this approach to quantitatively compare two systems: regenerating cell walls in plant protoplasts and cellulose nanofiber-pectin synthetic capsules. If native wall architectural features, such as cross-linking or cross-lamellar organization, are essential for mechanical properties, we expect the synthetic capsules would be unable to capture the mechanical properties of plant cell walls.

Wall regeneration has been studied in both *Arabidopsis thaliana* and *Nicotiana tabacum* (BY2) (39–41) cell lines. Cellulose architecture during wall regeneration has been visualized in protoplasts derived from Arabidopsis rosette leaves using advanced microscopy (39, 42, 43). Mechanical characterization of intact BY2 cells (44, 45) and Arabidopsis isolated cells showed that turgor pressure dominates cell stiffness (37, 46). However, the mechanical properties of the regenerating walls remain poorly quantified. We here focus on BY2 protoplasts because their rapid growth and high regenerative capacity in suspension culture make them ideal for monitoring dynamic changes in cell wall mechanics.

In this study, we integrate structural (Atomic Force Microscopy, AFM and Electronic Transmission Microscopy, TEM), compositional and mechanical (viscoelastic) analyses to directly compare synthetic primary cell wall analogues with regenerating walls of BY2 protoplasts. At selected stages during regeneration, we assess surface structure via AFM scanning, wall compositions with polysaccharide staining and mass spectroscopy, and mechanical properties using compression assays. Our results show that synthetic layered cellulose nanofiber-pectin walls can recapitulate viscoelastic characteristics of regenerating primary cell walls, suggesting that these two components are sufficient to confer key mechanical properties.

## 2. Results

### Regenerating a cell wall versus building a synthetic wall

To investigate de novo cell wall formation, we first generated wall-less plant cells, protoplasts, from BY2 *Nicotiana tabacum* cells in liquid culture. Protoplasts were obtained by a combination of cell wall degradation and hypo-osmotic shock (details in methods). Protoplasts were cultured in primary cell wall regeneration medium for 5 consecutive days (Scheme 1A and 1C). On day 0, immediately after protoplasting, protoplasts were spherical, averaging 38.3 ± 3.2 μm in diameter (Supplementary Fig. 3; Scheme 1C). After 5 days in the regeneration medium, most cells lost their spherical shape (Scheme 1C), indicating that a new cell wall had formed. By this time, their average diameter increased to around 55 ± 13 μm (Supplementary Fig. 3).

Synthetic cell wall analogues were prepared via the bottom-up layer-by-layer (Lbl) assembly protocol, involving the alternate adsorption steps of pectin and cationically modified cellulose nanofibers (CNFs) onto calcium carbonate (CaCO_3_) microparticle templates, as schematically illustrated in Scheme 1B. Each bilayer (bL) comprises one layer of pectin followed by one layer of CNFs and the total number of bilayers in the walls can be varied. We supposed that cell wall thickening occurred during the wall regeneration in BY2 cells, hence, synthetic analogues were prepared with 3 to 6 bilayers. These multilayered structures constitute polyelectrolyte multilayers, also referred to as polyelectrolyte complexes, which form spontaneously when oppositely charged macromolecules interact (47). The pectin carried a negative charge due to its carboxyl groups, while the surfaces of the CNFs were rendered positively charged through the covalent introduction of quaternary ammonium moieties. The assembly of polyelectrolyte complexes is primarily driven by the entropic release of counterions and water molecules upon complex formation (48). The cationic charge on the CNFs was needed in the layer-by-layer assembly. Following Lbl assembly, the CaCO_3_ template was dissolved using citric acid solution, and the residual salts were removed via thorough washing with MilliQ-water, finally yielding a core-shell structure with a water-filled interior (Scheme 1B and 1D). The final size of the 6 bL capsules after core removal was 15.8 ± 2.7 μm in diameter (Supplementary Fig.3), about half that of protoplasts. Table 1 provides a summary comparison of the regenerating and synthetic walls.

**Table 1.** Comparison between structural and mechanical properties between the synthetic and regenerated cell walls.

| **Property** | **Synthetic wall**<br>**(LbI capsules)** | **Regenerated wall**<br>**(BY2 cells)** |
| --- | --- | --- |
| **Diameter** | $\approx 15.8 \pm 2.7 \mu\text{m}$ (6bL) | $\approx 55 \pm 13 \mu\text{m}$ (day 5) |
| **Morphology** | Spherical | Spherical (day0) → elongated (day5) |
| **Wall assembly** | Layer-by-layer adsorption | Natural synthesis, wall secretion |
| **Wall composition** | Cellulose (CNFs),<br>Pectin,<br>Xyloglucan,<br>Mannan | Cellulose microfibrils ,<br>Pectin,<br>Xyloglucan,<br>Mannan |
| **Covalent cross-links**<br>**between wall**<br>**polymers** | No (electrostatic, hydrogen bonding,<br>van der Waals interactions). | Yes, in addition to non-covalent<br>cross-links, hydrogen bonding,<br>van der Waals, electrostatic<br>interactions. |
| **Wall architecture** | Multilayered wall, random CNF<br>orientation in layers | cross-lamellate cellulose<br>microfibrils organization in layers |
| **Cellulose**<br>**distribution in wall** | Homogenous | Heterogenous |
| **Turgor** | No turgor | Yes or No (when cells are<br>plasmolysed) |
| **Growth** | No | Yes |
| **Wall thickness** | $\approx 70 - 180 \text{ nm}$ (3bL – 5 bL) | $120 - 250 \text{ nm}$ (3 – 5 days) |

### Nanoscale structural comparisons

The nanoscale structure of the outermost cell wall layer in regenerating protoplasts at defined time points (post-regeneration) was investigated with AFM. The same measurements were performed on hydrated synthetic analogues with walls of varying numbers of bilayers (Fig. 1). Progressive changes in surface ultrastructure were observed during wall regeneration in BY2 cells. At day 1, cellulose microfibrils were not discernible at the protoplast cell surface (Fig. 1 *A*), whereas by day 2, nascent microfibrillar structures became evident (arrows in Fig. 1 *B*), and became more numerous by day 3 and day 5 (Fig 1 C and D). Given that cellulose microfibrils are synthesized and extruded by cellulose synthase complexes embedded in the plasma membrane, their presence in the outermost cell wall layer was anticipated from the early stages (day 1) of wall regeneration. Also, the outermost cell wall layer was expected to be similar at the different time points; 1, 2, 3 and 5 days. Cellulose staining with Calcofluor white was apparent from day 1 onwards (Fig 3H-K), indicating there is no delay in cellulose synthesis. The difference and delayed detectability of cellulose microfibrils with AFM suggest that the final and robust structure appears between day 2 and 3 of cell wall regeneration. AFM scanning reveals surface features by interacting mechanically with the structure; an unstable structure might be difficult to detect by this technique, i.e. at early stages, the wall organization might be unstable. Indeed, cellulose reportedly exhibits highly dynamic behavior at early stages of wall regeneration because of diffusion of short fibrils prior to densification of the network of elongating microfibrils (43).

**Fig. 1.**
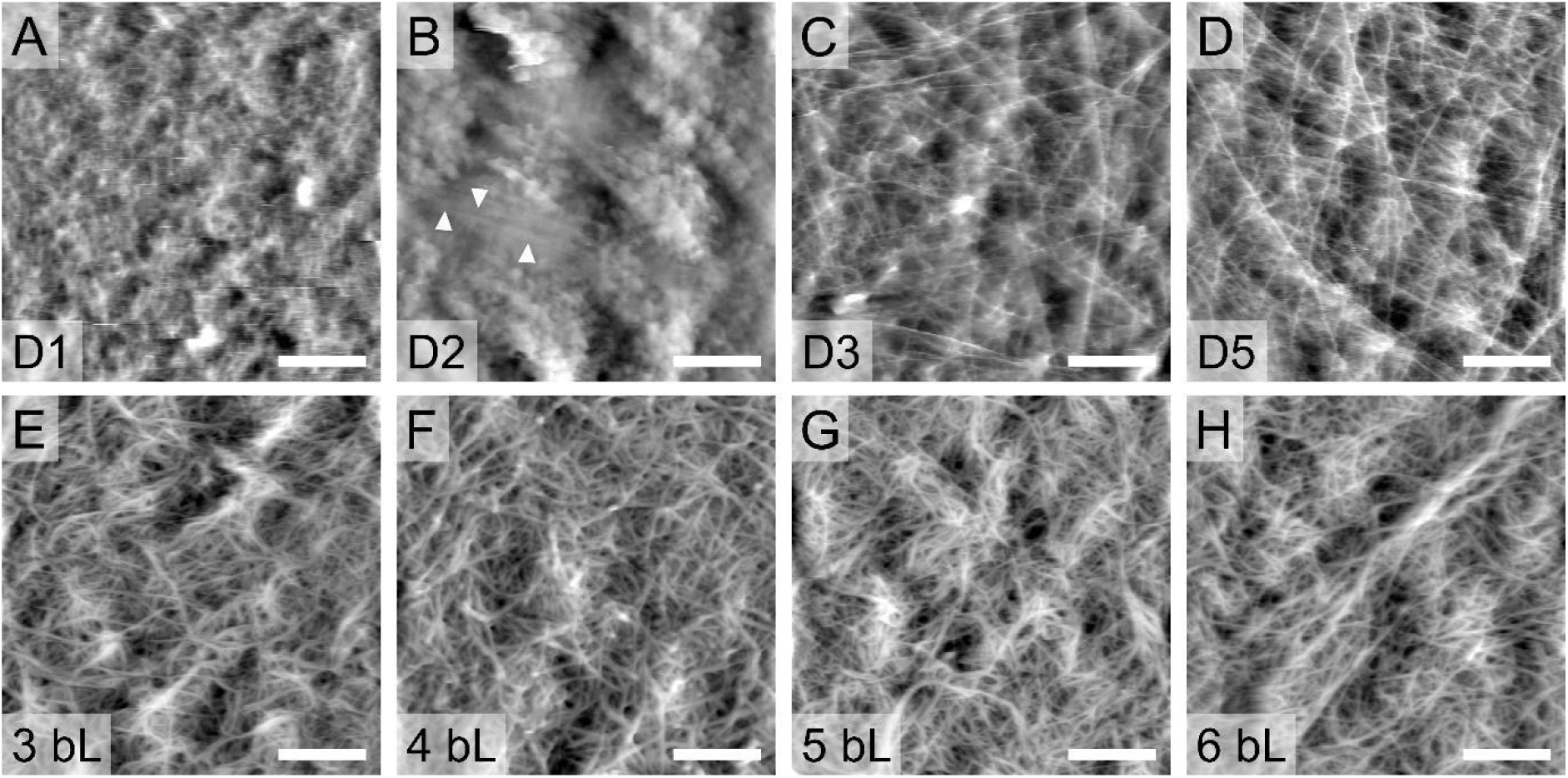
AFM of BY2 protoplasts and Lbl capsule surfaces. Regenerating protoplast (A) day 1, (B) day 2, (C) day 3, and (D) day 5 post-harvest. Synthetic wall analogues assembled with (E) 3 bL, (F) 4 bL, (G) 5 bL and (H) 6 bL of pectin and CNFs. All panels show Height Sensor AFM channel (first order plane fit, fifth order flatten). Arrows in B show fibers. Scale bars = 500 nm.

From day 3 onward, the characteristic cross-lamellate architecture of cellulose microfibrils in the primary cell wall became apparent and this distinct fibrillar morphology persisted through day 5, as shown in Fig. 1 *C* and *D*. From the micrographs, we observed that the integrated cellulose microfibrils organization was random.

Fig. 1 *E* - *H* presents AFM micrographs of the outermost surface layer of the synthetic analogues with walls that were composed of 3, 4, 5 or 6 bLs. The synthetic walls contained randomly oriented fibers, irrespective of the number of bilayers. The cross-lamellate structure was not replicated in the synthetic analogues. Additionally, compared to the cellulose microfibrils observed in the regenerated walls (Fig. 1 *C-D*), the cellulose nanofibers in the synthetic analogues exhibited greater bundling. CNFs displayed a higher degree of curvature and lacked the extended, linear morphology of the cellulose microfibrils in the native cross-lamellate arrays (Fig. 1*C-D*).

Wall thicknesses for regenerated and synthetic walls were evaluated from transmission electron microscopy (TEM) cross-sections (Fig. 2). The synthetic constructs exhibited a layered architecture (Fig 2D), which was further corroborated using polarized optical microscopy (POM) (Fig. 2E), where the appearance of Maltese cross patterns in the 5-bilayer samples provided additional confirmation of birefringence associated with layered ordering (POM of 3bL, 4bL and 5BL in Supplementary Fig. 2 in SI). The overall thickness of the synthetic walls increased with the number of bilayers: 71 ± 20 nm for 3bL, 124 ± 31 nm for 4bL, and 178 ± 42 nm for 5bL constructs (Fig. 2A). These thicknesses were matched by the thicknesses of regenerated BY2 cell walls (Fig 2A-B). The thickness of the regenerated walls increased with time: 130 ± 45 nm for day 3 and 204 ± 102 nm for day 5. In both systems, material accumulation led to wall thickening.

**Fig. 2.**
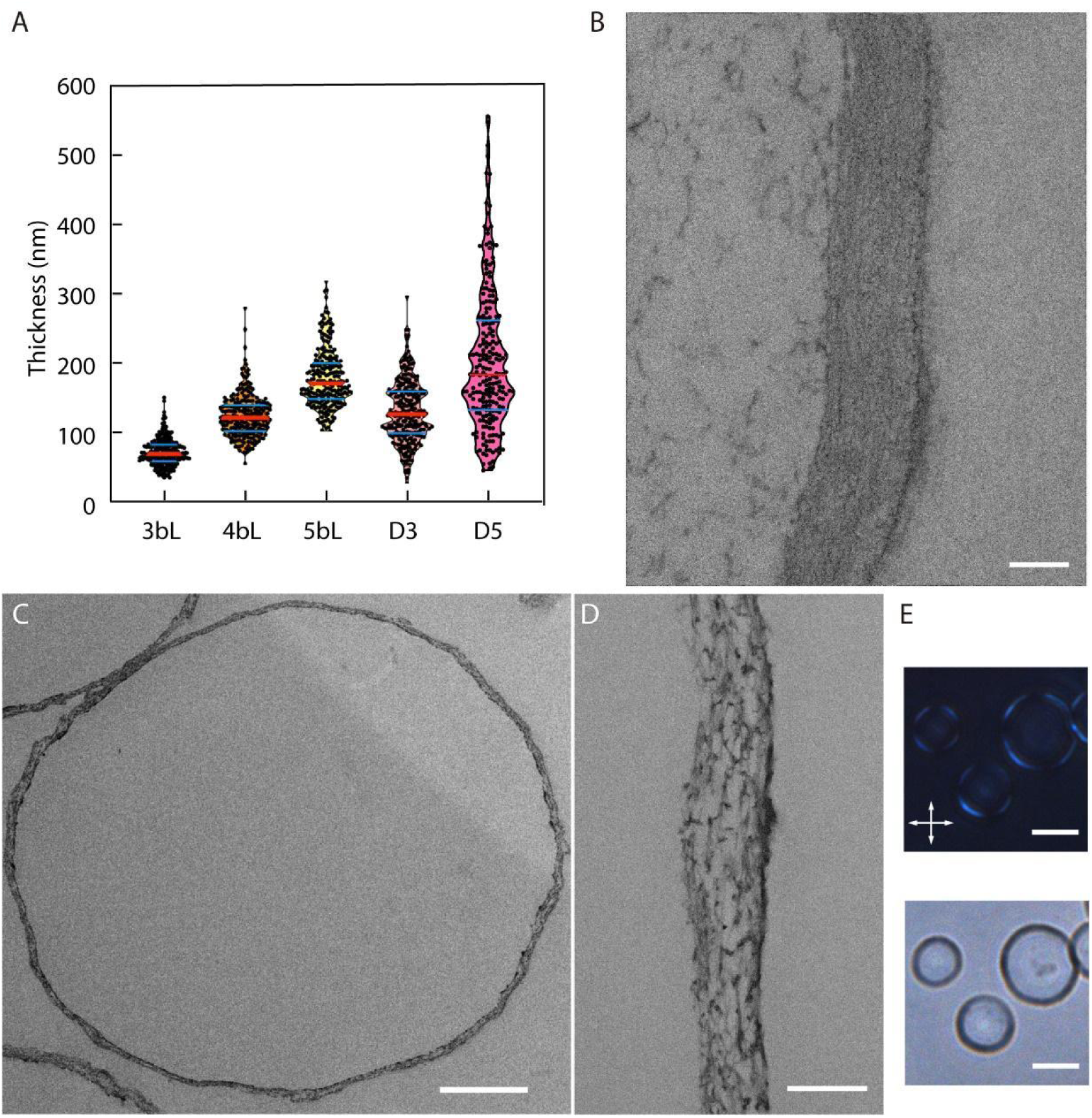
(A) Thicknesses measured from TEM of cross-sections of synthetic capsules (3bL,4bL and 5bL) and regenerated walls (day 3 and 5). (B) Cross-section of a regenerated wall (day 5). (C) Cross-section of a synthetic wall with 5 bilayers (3 and 4 bL TEM images are in SI). (D) close-up on the synthetic wall in (C). (E) Polarized optical microscopy (Top) and Brightfield (Bottom) images of a synthetic wall with 5 bilayers. Scale bars = 200 nm (B), 2 µm (C), 200nm (D), and 10 µm (E).

**Fig. 3.**
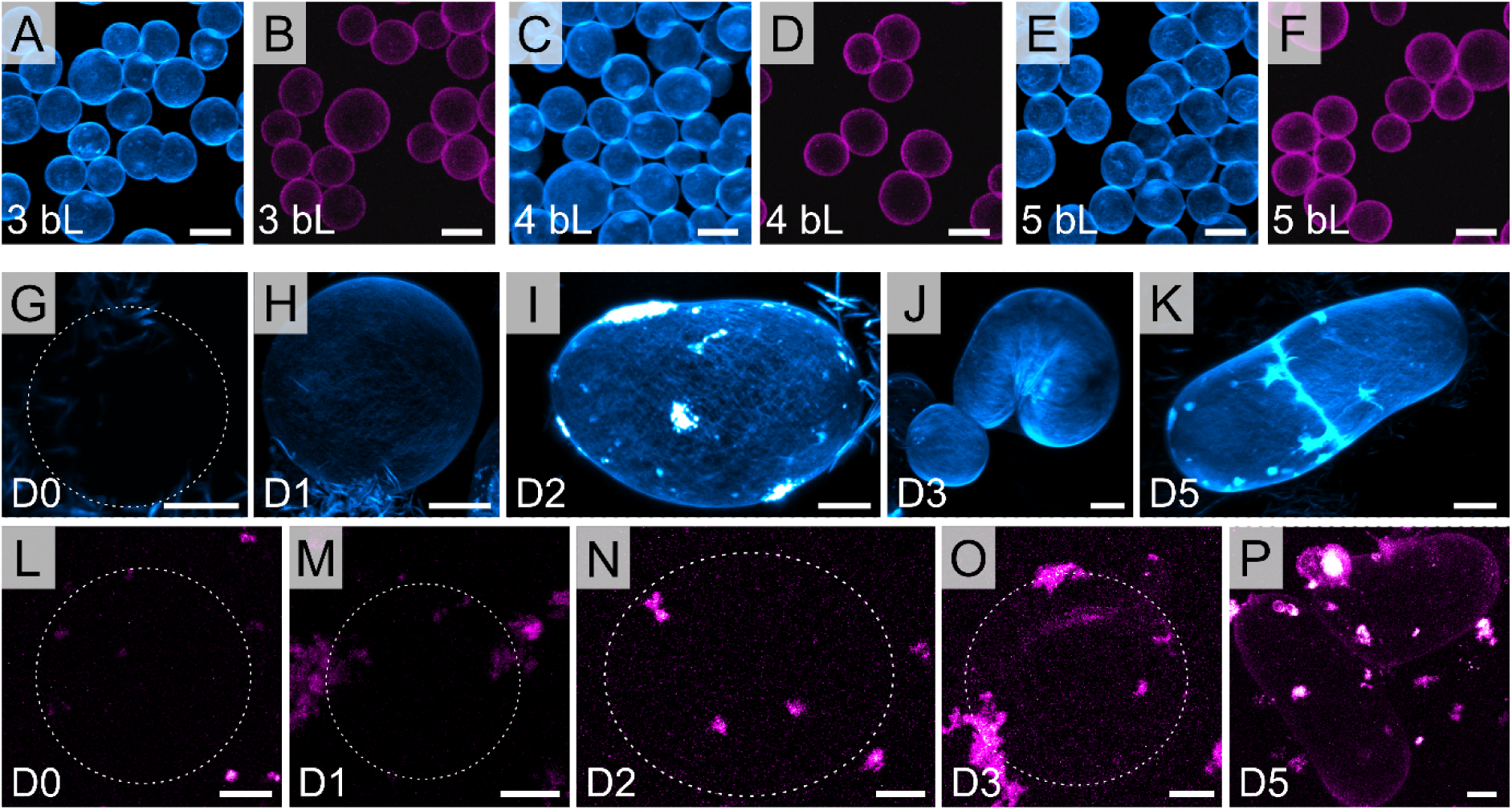
CLSM images of synthetic analogues (upper row) with 3bL (A, B), 4bL (C, D), or 5bL (E, F) and regenerating BY2 protoplasts (middle and bottom rows) at day 0 (G, L), day 1 (H, M), day 2 (I, N), day 3 (J, O) and day 5 (K, P) of cell wall regeneration. The cellulose was stained with Calcofluor white (CFW, cyan) and pectin with propidium iodide (PI, magenta) both in synthetic analogues (A-F) and the regenerated cell walls (G-K). Dotted lines show cell outlines in images where the cell is not clearly visible. Scale bars = 10 µm.

### Comparison of wall composition

Cellulose and pectin in the regenerated and synthetic cell walls were stained with Calcofluor White (CFW) and propidium iodide (PI), respectively, and the resulting confocal laser scanning microscopy (CLSM) images are presented in Fig. 3. As previously mentioned, BY2 protoplasts and the synthetic analogues exhibited different size ranges. On day 0, immediately after protoplasting, the protoplasts showed neither CFW nor PI staining, confirming that the cell wall had been largely removed during the process. Staining confirmed the presence of the two main polysaccharides, cellulose and pectin, in the synthetic and regenerated walls (from day 1 to day 5). Cellulose staining was consistently homogeneous across all investigated synthetic wall bL numbers. By contrast, the regenerating walls of BY-2 cells showed a less uniform pattern, with brighter regions indicating areas of higher cellulose concentration. The intensity of cellulose staining increased from day 1 to day 5 of wall regeneration, suggesting that the cellulose network became more established and denser as regeneration progressed. This pattern may reflect a dynamic organization of the cellulose network, as previously described by Huh et al. (43). PI binds to -COOH groups and we observed that the fluorescence PI signal was detectable after day 3 for the native regenerating walls, indicating that either pectins were not synthetized in the early stages of regeneration (before day 3) or that there was change in the pectin’s molecular composition at day 3. The pectin employed in the fabrication of the synthetic analogues was apple-derived and exhibited a high degree of esterification (70%).

Gas chromatography–mass spectrometry (GC-MS) analysis of the walls was performed to gain detailed insight into their biochemical composition. While imaging and staining techniques reveal structural and spatial information, GC-MS provides a sensitive and quantitative profile of part of the wall’s molecular components. This allows for the identification of specific monosaccharides, offering a more comprehensive understanding of how the wall composition differs between regenerating and synthetic systems. For the synthetic and regenerated walls (at day 5), the compositional characteristics of monomers are provided in Table S1 of the Supplementary Information. The synthetic walls exhibited higher monosaccharide contents compared to the regenerated wall on day 5. This difference is particularly pronounced for uronic acids (galacturonic and glucuronic acids), as well as for glucose and galactose. Overall, the composition highlighted a marked heterogeneity between the two types of walls. Although the relative abundances of monosaccharides type differed substantially between the synthetic and regenerated walls, the types of monosaccharides present were largely similar between the two systems.

### The wall of regenerating protoplasts stiffens as wall thickens

As the regenerating cell wall thickens, it is expected to gain mechanical strength. The mechanical properties of cells at different stages of wall regeneration were evaluated using a single-cell rheometer composed of two parallel plates. A single cell was captured between two custom-made microplates: one rigid and the other flexible, with a calibrated stiffness, functioning like a spring to apply controlled forces to the cell. Uniaxial oscillatory compression was then applied with the flexible plate. During this process, both the deformation of the sample and the applied force were recorded. This approach enabled us to quantify the overall viscoelastic response of the cell, capturing both the storage (K’) and loss (K’’) stiffness as a function of the number of days of wall regeneration and turgor modulation (details in mat and methods). The storage stiffness K’ relates to the elastic response of the sample, while the loss stiffness relates to the energy dissipation by viscous effects within the sample. An increase in storage stiffness was observed during cell wall regeneration (Fig. 4C): K’ increased from 20 ± 11 nN/µm (day 1) on average to 760 ± 195 nN/µm (day 5). Thus, storage stiffness increased by approximately three orders of magnitude over five days. The loss stiffness K’’ increased over the five-day time course, rising by approximately two orders of magnitude from 0.1 nN/µm on day 0 to 10 nN/µm on day 5. Unlike K’, a plateau was observed around day 3 in control condition, suggesting that the loss stiffness stabilizes faster than the elastic stiffness. Although the cultures could be maintained beyond five days, eventually leading to full regeneration of the callus tissue, mechanical measurements were limited to the first five days. Beyond this period, the onset of cell division introduced multicellularity and structural heterogeneity, thereby interfering with accurate mechanical measurements.

**Fig. 4.**
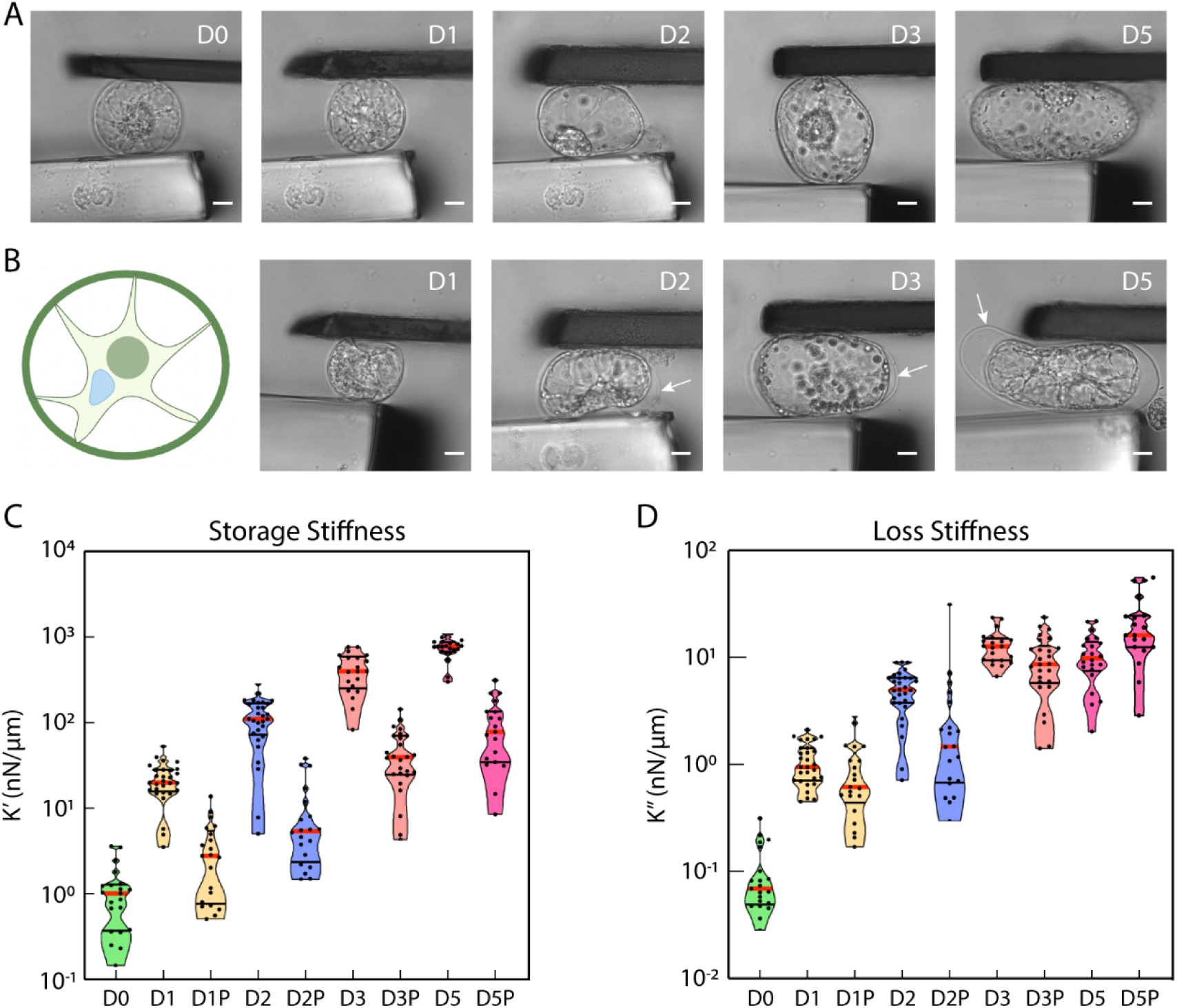
Storage and loss stiffnesses for regenerating BY2 cells. Light microscopy images of cells at each day of regeneration, in both control (A) and plasmolysed (B) conditions, Scale bars = 10 µm. White arrows indicate the cell wall in plasmolysed condition. (C) storage stiffness (K’) and (D) loss stiffness (K’’) are plotted as a function of the days of cell wall regeneration.

To assess the mechanical properties of the wall in the absence of turgor pressure, viscoelastic measurements were performed in culture medium supplemented with mannitol to impose hyperosmotic stress. The results (Fig. 4) showed that single-cell mechanical assays are highly sensitive to turgor pressure, which is in line with previous studies (37, 45, 46, 49). At each time point, an equal concentration of mannitol was added to the medium to induce plasmolysis, evidenced by detachment of the plasma membrane from the cell wall (arrows, Fig. 4B). At all experimental time points (1, 2, 3, 5 days), the measured storage stiffness (K’) (Fig. 4C) decreased by approximately one order of magnitude upon plasmolysis. K’ measured under plasmolyzed conditions, reflecting primarily the mechanical properties of the cell wall, increased by about two orders of magnitude over the five-day regeneration period. This result supports the conclusion that the plant cell wall stiffness increases as the wall thickens.

Our findings demonstrate that turgor pressure is the dominant contributor to the overall storage stiffness of the cell. Quantitative analysis revealed that the cell wall accounts for approximately 10% of the overall elasticity, with the remainder primarily attributable to internal turgor pressure. The loss stiffness (K’’) was minimally affected by hyperosmotic shock (Fig. 4D), as plasmolyzed cells exhibited a loss stiffness equivalent to turgid cells. This indicates that the rise of the viscous behavior is primarily due to the regenerating cell wall, which contributed approximately 80% of the measured loss stiffness, with limited input from turgor pressure.

### Synthetic capsules recapitulate the mechanical behavior of regenerating cell walls

To compare mechanical properties, we measured both the synthetic walls and regenerating plant cell walls using the same microplate-based compression device in aqueous conditions. Oscillatory compression tests revealed that both systems exhibit similar viscoelastic signatures: across the tested frequency range, storage stiffness (K’) consistently exceeded loss stiffness (K’’), indicating that the mechanical behavior is predominantly elastic. In both cases, storage stiffness followed a power-law dependence on frequency, *K*^’^ = *K*^’^_0_ *f*^α^, with comparable exponents α= 0.06 ± 0.02 for the synthetic wall and α= 0.12 ± 0.07 for the regenerated wall at day 5 (Figure 5B-C), suggesting similar viscoelastic behavior.

The synthetic capsules also exhibited a monotonic increase in K’ and K’’ as the number of layers increased (Fig 5D-E). Since the thickness of the synthetic wall increases with the number of layers (see Fig. 2), the observed rise in elasticity reflects the accumulation of wall material. Notably, the measured values for the synthetic capsules fall within the same range as those of regenerated plant cell walls at day 5 (Fig. 5D), highlighting the synthetic wall’s ability to mimic both the elastic and dissipative characteristics of native walls.

**Fig. 5.**
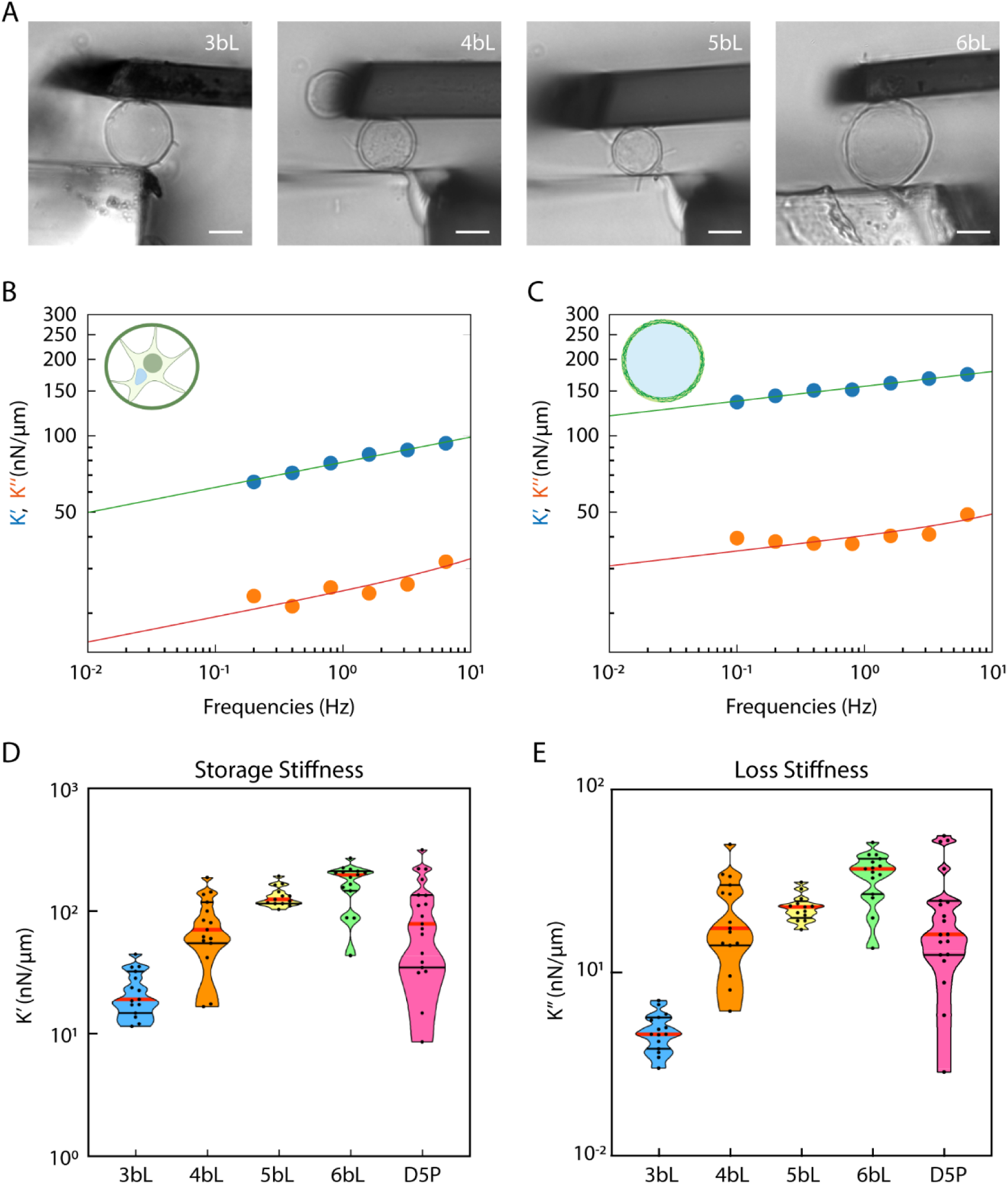
Storage stiffness and loss stiffness of synthetic wall analogues and day 5 plasmolysed cells. Light microscopy images of the synthetic analogues are shown in (A) (scale bars = 10 µm). Storage stiffness (K’, green) and loss stiffness (K’’, blue) as a function of frequency for day 5 plasmolysed cell (B) and synthetic wall analogues (C). Storage stiffness (K’, D) and loss stiffness (K’’, E) are plotted as a function of thickness for synthetic analogues composed of 6 bilayers (6 bL, n = 15), 5 bilayers (5 bL, n = 13), 4 bilayers (4 bL, n = 15), and 3 bilayers (3 bL, n = 15).

### Wall material quantity controls stiffness evolution in both systems

The smaller radius of synthetic capsules compared to regenerated cells will increase the value of K’. K’ also increases with wall thickness. To the compare the two systems, we therefore estimated the effective Young’s modulus *E*^’^ of the walls from the classical thin-shell theory (50): *E*^’^ = *K*^’^*R*/*h*^2^, where R is the shell radius, and h is the wall thickness. For both systems, the resulting effective Young’s modulus was independent of the wall thickness (Fig. 6). This reveals that the increase in stiffness reported in the previous sections is due to the material accumulation in the wall through the addition of layers for the synthetic system or through regeneration days for the plant wall. The effective Young’s moduli fall in the same range of values for both systems.

**Fig. 6.**
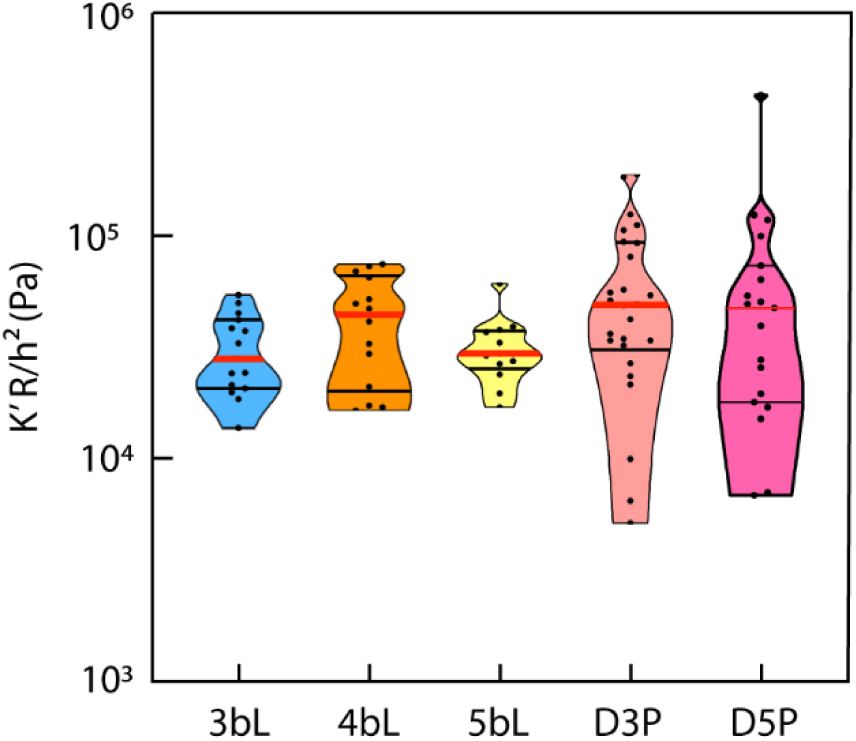
Effective elastic modulus (E’) as a function of thickness. E’=*K’·R/ h^2^* is plotted for synthetic wall analogues composed of 6 bilayers (6 bL, n = 15), 5 bilayers (5 bL, n = 13), 4 bilayers (4 bL, n = 15), and 3 bilayers (3 bL, n = 15), as well as for D3 and D5 plasmolysed cells (n = 26 and n = 19, respectively).

Taken together, these results demonstrate that the synthetic cellulose-pectin capsules not only replicate the magnitude of elastic and loss stiffness of regenerating plant cell walls but also recapitulate the scaling behavior with wall thickness and frequency. Thus, synthetic walls capture the essential mechanical features of native primary cell walls and provide a physical model for dissecting plant wall mechanics.

## 3. Discussion

Plant cell walls are complex, with highly oriented cellulose microfibrils embedded within a heterogeneous matrix of polysaccharides that are constantly being remodeled. By contrast, the synthetic cell wall analogues offer a simplified and well-controlled platform to test specific hypotheses. These analogues consist of stratified, alternating layers of pectin and cellulose nanofibers. They are structurally static and lack remodeling capacity. Our AFM analysis indicated that cellulose nanofibers in the synthetic wall are more curved and bundled and less extended compared to the cellulose microfibril cross-lamellate structure observed in regenerated walls on day 3 and 5. Despites these differences in nanoscale architecture, we show that synthetic capsules capture key macroscopic mechanical properties of regenerating cell walls, including storage and loss stiffnesses. Such observations align with findings in other biomimetic materials. For example, natural nacre (mother-of-pearl) exhibits remarkable strength and toughness thanks to a highly ordered “brick-and-mortar” architecture of aragonite platelets and softer organic layers (51). Yet, synthetic alumina composites produced by freeze casting can match nacre’s mechanical performance despite a less ordered internal structure (52). Similarly, biomimetic spider silk crosslinked with multi-arm polyethylene glycol attains mechanical properties close to natural silk, including tensile strength and toughness despite distinct molecular and fiber arrangement differences (53, 54).

Young’s modulus of plant cell walls ranges from 0.1–10 MPa in meristematic tissues (18) (55) and ∼0.3 MPa for turgid BY-2 cells (46). In contrast, the effective Young’s modulus on plasmolyzed BY-2 cells with regenerated walls measured in this study is much lower (∼0.06 MPa at day 5). The lower values reported here might indicate that the regenerated wall had not achieved full mechanical maturity at day 5. Stiffness measured on turgid cells (0.76 N/m) is also lower than the stiffness measured previously on intact BY2 cells with Cellular Force Microscopy (4–8 N/m, (45)), indicating that turgor pressure is lower in cells with 5 days regenerated walls or that the cell wall is weaker than in intact BY2 cells.

In their construction, the synthetic wall analogues required electrostatic interactions, in particular, cationic modification of CNFs, for layer-by-layer assembly. By comparison, cellulose microfibrils in living plant walls are not positively charged. Nevertheless, negatively charged pectin in native walls associates with cationic partners, for example through homogalacturonan forming pectate coacervates with positively charged extensin proteins (56). This demonstrates that similar mechanical behavior can emerge from different molecular interactions.

In primary cell walls, various types of molecular interactions and bonds between pectin and cellulose (57, 58) and between cellulose and xyloglucan (27) have been reported (Table 1) and certain bonds have been linked to wall strength (27, 59). To which degree such molecular interactions/bonds spontaneously form in our synthetic capsule walls remain uncertain. Nonetheless, the mechanical properties of regenerating walls fall in the same range as those of intact plant cell walls and match those of synthetic capsules. Our findings thus challenge the notion that a specific type of molecular interaction (or bond) between pectin-cellulose and xyloglucan-cellulose is essential for wall strength. This interpretation would be consistent with recent work showing that wall mechanics are primarily sustained by interactions between cellulose fibrils (60).

TEM imaging showed that synthetic walls exhibited a stratified and uniform thickness throughout their structure, whereas native walls displayed variable layering and thicknesses. Lipowczan et al. (61) suggested that plant cell walls are organized in multiple layers of varying thickness, creating elastic strain gradients that reflect both deposition history and remodeling, thereby guiding growth and morphogenesis. While the TEM sample preparation might have generated wall thickness variation in the native wall, the thickness variability identified in our measurements might also reflect wall shaping by local developmental cues and remodeling during wall regeneration. By contrast, synthetic walls exhibit a stratified but uniform thickness, reflecting the static and controlled nature of these artificial systems. Nonetheless, at the global cell scale, the layered architecture of pectin and cellulose appears to be sufficient to match regenerating walls mechanical properties.

Thickness-dependent mechanics are well established for polyelectrolyte multilayer microcapsules. When tested in aqueous environments, micro-capsule stiffness scales with wall thickness squared (50). Our quantification shows that the regenerating walls and the synthetic capsules exhibit increasing storage and loss stiffnesses as their walls thicken, showing comparable thickness-dependent mechanical trends. This highlights how the overall amount of material, rather than specific architectural details, governs mechanical behavior in the early days of regeneration.

Pectin state (unsubstituted or methyl-esterified) is expected to change during wall regeneration (62). In confocal microscopy, our imaging showed that the PI staining became significant around day 3 after wall regeneration started. Because PI binds to demethylestrified pectin (63), methylesterified pectins might be secreted before day 3, and our results suggest that from day 3 onwards they undergo a change in their substitution state, becoming less methylated. AFM imaging highlighted the emergence of a well-organized structure in the cell wall beginning on day 3. This timing coincides with the PI staining becoming significant and with the plateau observed in the loss stiffness, which stabilizes earlier than the storage stiffness (Fig. 4). This temporal correlation suggests that the organized structure with a change in pectin state may be sufficient to confer dissipative mechanical properties to the cell wall. In contrast, the full development of its storage capacity might require further structural reinforcement, potentially mediated by the crosslinking polymer networks (43).

Confocal microscopy revealed that the distribution of polymers such as cellulose and pectin is more homogeneous in synthetic capsules compared to native cell walls. The spatial heterogeneity in polymer density found in regenerating walls might reflect the dynamic regulation of wall deposition as highlighted in other studies (39, 43). It is also proposed that local polymer organization influences cell morphogenesis and mechanics (64, 65). In addition, non-uniform organization may influence mechanical consequences, as shown by Peaucelle et al. (18), who suggested that local pectin de-methylesterification modulates wall elasticity, triggering organ formation. In contrast, the homogeneous polymer distribution in synthetic capsules reflects the static and controlled nature of these artificial systems. Similar observations have been reported in other capsule types, including alginate–PLL–alginate microcapsules (66) and microcapsules composed of natural polymers (67), both of which display uniformly distributed polymers.

## Conclusion

Plant cell growth depends on the mechanics of the primary wall, which reflect its composition, organization, and remodeling. Comparing regenerating BY2 walls with simplified cellulose–pectin capsules shows that native architectures are not required to reproduce key mechanical behaviors: both systems display thickness-dependent increases in stiffness and viscous dissipation despite distinct nanoscale organizations. This suggests that layered cellulose and pectin are sufficient to capture essential viscoelastic properties.

More broadly, the structural complexity of native walls likely serves dynamic functions at long time scales —such as anisotropic growth, remodeling, and stress adaptation—rather than being indispensable for static mechanical integrity. Synthetic capsules, therefore provide a minimal yet powerful model to probe how additional polymers and remodeling processes shape wall mechanics.

## Materials and Methods

### 4.1 Materials

#### CNF sample preparation

Never-dried softwood pulp (Nordic Seffle AB, Sweden) were used in reaction with glycidyltrimethylammonium chloride, followed by mechanical disintegration, to generate the cationic (quaternary ammonium groups) CNFs, using a well-documented protocol as described in detail earlier (68). The cationic charge density was (1.2 mmol g^−1^ fiber) for the CNF (conductometric titration, (69)).

### 4.2 BY2 cell culture and protoplasting

BY-2 cells were cultured in BY-2 medium containing 4.67 g/L Murashige and Skoog (MS) basal salt mix without vitamins, 30 g/L sucrose, 0.2 g/L KH_2_PO_4_ (monobasic potassium phosphate), and 0.1 g/L myo-inositol. The pH was adjusted to 6.0 with 1M KOH prior to autoclaving. After sterilization, 1 mL/L thiamine (0.1 g/100 mL) and 200 µL/L of a 10 mg/mL 2,4-dichlorophenoxyacetic acid (2,4-D) stock solution were added aseptically. For callus culture, the medium was solidified with 7 g/L plant agar. Liquid cultures were initiated by transferring a small piece of 3 weeks old callus into 100 mL of liquid BY-2 medium and incubated at 25 °C with shaking at 130 rpm for one week to allow cell proliferation. After one week, 10 mL of the cell suspension were collected and centrifuged at 1200 rpm for 3 minutes. The resulting pellet was resuspended in 10 mL of enzymatic solution for protoplast isolation. This solution, prepared in BY-2 medium supplemented with 0.4 M mannitol and thiamine (without hormones), contained 850 mg cellulase R10, 850 mg cellulysin, and 20 mg pectolyase Y23 per 50 mL. The cell suspension was incubated for 4 hours at 60 rpm, room temperature, protected from light with aluminum foil.

Following enzymatic digestion, the suspension was centrifuged at 1200 rpm for 3 minutes, and the supernatant was discarded. The pellet was rinsed twice with BY2 medium with 0.4 M mannitol and thiamine, each rinse followed by centrifugation at 1200 rpm for 3 minutes and removal of the supernatant. A final wash was performed with BY2 medium supplemented with 0.2 M mannitol and thiamine. The protoplast suspension was then left to rest for 10 minutes in the dark and filtered through a 45 µm nylon mesh (Fisherbrand) to eliminate debris. After filtration, the suspension was centrifuged again at 1200 rpm for 3 minutes, and the supernatant was carefully removed. Protoplasts were finally resuspended in BY-2 medium supplemented with 0.2 M mannitol, thiamine, 200 µL/L of a 10 mg/mL 2,4-dichlorophenoxyacetic acid (2,4-D) stock solution, and 1 mL/L of a 1 mg/mL 6-benzylaminopurine (BAP) stock solution. The protoplast culture was incubated in the dark at 25 °C for 5 days to allow cell wall regeneration.

### 4.3 Layer-by-layer assembly of synthetic analogues

Apple pectin (0.1 wt%, Sigma) was dissolved in 100 mM NaCl and stirred overnight at room temperature. The CNF (0.05 wt%) was diluted with MilliQ-water and stirred overnight. The pH was adjusted to 7.5 ± 0.2 for both CNF and pectin using 0.1 M NaOH or HCl. The pectin solution was filtered through a 0.8 μm syringe filter (Corning, Germany). The CNF suspension were sonicated for 60 seconds (Sonics Vibra-Cell, 750 W, 80% amplitude, ½″ tip) and then centrifugated for 10 min and the supernatant was collected (sediment discarded). NaCl was then added to the CNF suspension to obtain a 100 mM NaCl content.

Microcapsules were prepared by sequential deposition of apple pectin and cellulose nanofibrils (CNF) onto CaCO_3_ microparticles using a layer-by-layer (Lbl) approach, as described previously (34, 70, 71). CaCO_3_ particles were prepared as described earlier (32) (72). An amount of 25 mg of dry CaCO_3_ particles were dispersed in 4 mL of 100 mM NaCl and sonicated at 40% amplitude for 10 seconds (Sonics Vibra-Cell, 750 W, ½″ tip). The particles were then collected by centrifugation at 5000 rpm for 1 minute. The CaCO_3_ was resuspended in 2 mL of the 0.1 wt% pectin solution (100 mM NaCl, pH 7.5 ± 0.2) and vortexed for 18 minutes (Vortex Genie, speed 10) to allow pectin to adsorb on the CaCO_3_ particles. After centrifugation, the supernatant was removed, and the particles were washed three times by resuspending in 2 mL of 100 mM NaCl solution, vortexing for 1 minute, and centrifuging under the same conditions. Next, the CNF layer was deposited by incubating the washed particles in 2 mL of the 0.05 wt% CNF suspension (100 mM NaCl, pH 7.5 ± 0.2) and vortexed for 18 minutes, followed by the same washing procedure. This deposition cycle was repeated to obtain the desired number of bilayers (3 to 6). Following Lbl assembly, the coated CaCO_3_ particles were rinsed with 100 mM NaCl for 1 time, then with MilliQ-water for 5 times and then dried at 50 °C overnight. Prior mechanical testing, dried particles were rehydrated in MilliQ water for 10-60 minutes and then incubated in 50 mM citric acid at least 1h to allow the CaCO_3_ core to dissolve. The capsules were then thoroughly washed with MilliQ water and stored at room temperature in water prior to further characterisation.

### 4.4. Characterisation

#### Atomic Force Microscopy

AFM imaging was performed as previously described in (73). Protoplasts were washed in BY-2 medium, pelleted by centrifugation, and a small volume of cells was deposited on silane-coated glass microscope slides and allowed to dry completely. At the time of scanning, the protoplasts were rehydrated in ddH2O and washed until no further debris was removed. The Lbl capsules were similarly rehydrated on glass slides at the time of AFM scanning. Capsule cores were removed as described prior and after washing, hydration was maintained with ddH2O throughout the scanning process. AFM topography images were captured on a Dimension Icon AFM (Bruker, CA, USA) using a calibrated Scanasyst-Fluid + probe (spring constant of 0.7 N/m, nominal tip radius of 2 nm; Bruker). 2 µm × 2 µm images were captured in QNM PeakForce Tapping mode (in fluid) at a resolution of 1,024 pixels/line. All images in figures are the Height Sensor AFM channel, first order plane fit, and fifth order flattened using Nanoscope Analysis V 2.0 (Bruker) software, to show details.

#### Transmission electron microscopy

BY-2 cells were fixed daily in a 1% (v/v) glutaraldehyde (GA) solution prepared in BY-2 medium supplemented with 0.2 M mannitol. The fixation was carried out overnight at 15 °C with gentle agitation. The following day, cells were centrifuged for 2 min at 1300 rpm, the supernatant was removed, and the pellet was resuspended in 500 µL of BY-2 medium containing 0.2 M mannitol.

The cell suspension was then transferred into sealed capsules, which were placed in Eppendorf tubes and centrifuged again for 2 min at 1300 rpm. After removal of the residual supernatant, cells were overlaid with 500 µL of 20% gelatin (Dr. Oetker) prepared in 0.2 M mannitol and maintained at ∼37–40 °C to remain liquid. Capsules were left at room temperature until the gelatin solidified completely. Additional fixation was done with 2.5% GA.

For the synthetic analogues, core-removed capsules in water were fixed with 2.5 wt% GA for 2h.

After fixation, all samples were transferred to EtOH absolute in steps (50, 70, 90 and ≥ 99.5 vol%) and embedded in epoxy (Epoxy embedding medium kit Sigma Aldrich) in polyethylene moulds (BEEM capsules). The resin was hardened at 45 °C (12 h), followed by 60 °C (24 h). Samples were cut with a Leica Ultramicrotome (EM FC7). The ultra-thin sections were stained with uranyl acetate (5 % in water) for 45 min. TEM micrographs were attained with a Thermo Scientific Talos L120C transmission electron microscope (acceleration voltage of 120 kV). The thickness of the wall was measured with ImageJ.

#### Polarised Optical microscopy (POM)

was attained using crossed linear polarizers in transmission mode and an Axio Vert.A1 Light Microscope (Carl Zeiss, Germany) equipped with a Zeiss AxioCam 305 color camera (Zeiss Zen 2.6 (blue edition) software), and a 63x/0.65 LD A-plan objective.

#### Confocal Laser Scanning microscopy (CLSM)

For each day of cell wall regeneration, protoplasts were stained with a final concentration of 0.02% (w/v) Calcofluor White Brightener 28 (CFW) from a 0.2 % stock solution and 0.01% propidium iodide (PI) prepared from a 1 mg/mL stock solution. Stained cells (50 µL) were mounted between a Superfrost microscope slide and a 1.5 thickness coverslip, separated by a spacer, and immersed in immersion oil (refractive index 1.334), compatible with the oil immersion objective used for imaging. Synthetic cell walls, with CaCO_3_ cores dissolved, were stained with 0.001% CFW (w/v) or 0.003% PI (w/v) and were similarly mounted for imaging.

Imaging of regenerated and synthetic walls was performed on Zeiss LSM 780 confocal microscope(s) equipped with a 63×/1.2 NA Olympus oil immersion objective, or a 63×/1.4 oil DIC M27 immersion objective (for the PI-stained synthetic cell walls). For dual-stained plant cells, single channels were acquired sequentially using “channel per track” mode to avoid cross-talk. The CFW signal was excited with the 405 nm laser at 2% power. PI was excited with the 543 nm laser at 27.9% power. Acquisition parameters were as follows: zoom 1.5, image size 512 × 512 pixels, pixel size 0.18 µm, and Z-step of 0.38 - 0.5 µm. CFW-stained synthetic walls were imaged with a 405 nm laser at 3% power (3bL, 4bL) or 2% power (5 bL), using the following parameters: zoom 1, image size 1024×1024 pixels, pixel size 0.13 µm x 0.13 µm and Z-step of 0.5 µm. PI-stained synthetic cell walls were imaged with a 514 nm laser at 25% power with the same acquisition parameters.

#### Gas chromatography–mass spectrometry

The monosaccharide composition of the cell wall was analysed. Three biological replicates, each consisting of approximately 10⁵–10⁶ BY2 cells (corresponding to ∼200 µg of dried cell wall), were subjected to three successive washes in an ascending ethanol gradient (50%, 80%, and 100%). A final wash was performed with acetone, after which the samples were dried overnight at room temperature. To remove residual starch, the dried material was incubated overnight at 37 °C with α-amylase from *Aspergillus oryzae* (Sigma, ref. A9857) in 200 µL of 100 mM ammonium acetate buffer (pH 5.15). After digestion, the supernatant was discarded and the pellet was washed twice with 500 µL of ammonium acetate buffer and one time ethanol 70% before drying.

For hydrolysis, the dried cell wall, previously weighed, was resuspended in 400 µL of freshly prepared trifluoroacetic acid (TFA, 2,5 M) and incubated at 120 °C for 1 h in 1.5 mL screw-cap tubes. This treatment specifically hydrolyzes the TFA-soluble polysaccharides (mainly hemicelluloses and pectins) into their constituent monosaccharides, while crystalline cellulose remains intact. After centrifugation (10 min), 10 μL of the supernatant was transferred and dried at room temperature using a speed-vacuum concentrator.

For derivation, 10 μL of 20 mg mL−1 methoxyamine in pyridine were added to the samples and the reaction was performed for 90 min at 28 °C under continuous shaking in an Eppendorf thermo-mixer. 50 μL of N-methyl-N-trimethylsilyl-tri uoroacetamide (MSTFA) (Aldrich 394866–10×1mL) were then added and the reaction continued for 30 min at 38 °C. After cooling, 45 μL were transferred to an Agilent vial for injection. Four hours after the end of derivatization, the whole sample series was injected in splitless mode. Five different standard mix were injected at the beginning and samples were randomized. An alkane mix (C10, C12, C15, C19, C22, C28, C32, C36) was injected for external retention index (RI) calibration. Injection volume was 1 μL. The instrument was an Agilent 7890A gas chromatograph coupled to an Agilent 5977B mass spectrometer.

Gas chromatography–mass spectrometry (GC-MS) conditions as well as data processing were performed as described in (74). In this study, peak areas were normalized to myo-inositol (internal standard) and to the dry weight of the samples.

#### Mechanical testing

Mechanical characterization of synthetic capsules and BY2 cells was performed with a parallel plate device as described previously in(37, 38). More precisely, we measured the storage stiffness (K’: elastic-like properties) and loss stiffness (K’’: dissipative, viscous-like features) at the whole cell scale by performing dynamical oscillations tests in the range of 0.1 to 6.4 Hz. The cell mechanical behavior as a function of the frequency can thus be represented by K’(f) and K’’(f).

In practice, a single cell or capsule was captured between two custom-made glass microplates under a bright-field microscope. One microplate is stiff compared to the sample while the other one is flexible with a calibrated stiffness k (1-147nN/μm). A sinusoidal displacement *D*(*t*) = *D*_0_*e* ^*i*ω*t*^ was imposed to the basis of the flexible microplate, leading thus to a sinusoidal compressive force applied on the cell. To determine the cell deformation, the displacement *d*(*t*) = *d*_0_ *e ^*i*ω*t*^*^+ ϕ^ of the flexible microplate, (with ϕ the phase shift between D and d) was measured through a S3979 position-sensitive detector (resolution∼200 nm). The instantaneous force can be calculated *F*(*t*) = *k*(*D*(*t*) − *d*(*t*)). Both *D*(*t*) and *d*(*t*) were well fitted by a sinus function. The elastic stiffness K’ and loss stiffness K’’ were then given by:

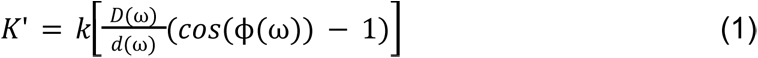

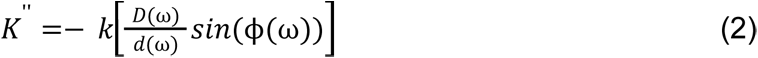

All results presented were performed for strains between 1-10% to avoid a nonlinear response due to too large strains. For all tested cells and synthetic walls, both K’ and K’’ exhibited a frequency dependence following a weak power-law behavior, described by:

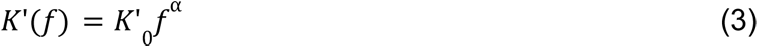

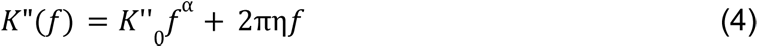

In the results section of the paper, the characteristic storage stiffness K’ and loss K’’ stiffness at the reference frequency f_0_ = 1Hz are reported (corresponding respectively to K’_0_ and K’’_0_ extracted from the power law fit after frequency sweep on each cell).

Compared with other techniques previously used to analyze plant cell mechanics, the parallel plates method allows us to measure a broad range of values of stiffness (unlike aspiration), and it provides an integrated cell-scale measurement of cell mechanics (unlike nano-indentation). Cells between the microplates were visualized under bright light illumination with a Plan Fluotar L 63×/0.70 objective and a Lumenera digital CCD camera (Infinity3, Lumenera, Canada). Vibration isolation was achieved by a TS-150 active antivibration table (HWL Scientific Instruments, Tübingen, Germany).

The storage (K’) and loss (K”) stiffnesses were measured on individual cells or individual bL capsules using the microplate system. For each experimental condition and for each day of cell wall regeneration, measurements were performed on a total of 20 to 30 cells, distributed over two independent experiments with 10 to 15 cells per experiment to ensure reproducibility.

#### Mannitol treatment for plasmolysis

Initially, turgid cells were placed in 10 mL of BY-2 medium supplemented with 0.2 M mannitol, corresponding to an osmolarity of approximately 425 mOsmol, measured with an osmometer. This condition represents the turgid state of the cells. To induce plasmolysis, 4 mL of BY-2 medium containing 0.7 M mannitol (osmolarity approximately 1020 mOsmol) was added, resulting in a final medium osmolarity of about 630 mOsmol. This hyperosmotic shock triggers cell plasmolysis. Mechanical measurements were conducted on plasmolyzed cells 20 minutes after the hyperosmotic treatment to allow stabilization of their physiological state.

## Acknowledgements

This study is supported by an HFSPO grant (grant number 2022-RG107). A.J.S, R.S and A. M. acknowledge VR application (grant number 2021-04761). We gratefully acknowledge Dr. Agnieszka Ziolkowska from Umeå Centre for Electron Microscopy for assisting with the TEM sample preparation. The study was partially supported by the Université Paris Cité, Idex ANR-18-IDEX-0001, funded by the French Government through its “Investments for the Future” program, and also by the projects “Mecha-Nuc” ANR-20CE13-0025-03 and “scEm-bryoMech” ANR-21-CE13-0046. The authors thank Liza A. Wilson for her advice and assistance with AFM imaging.

---

**Scheme 1.**
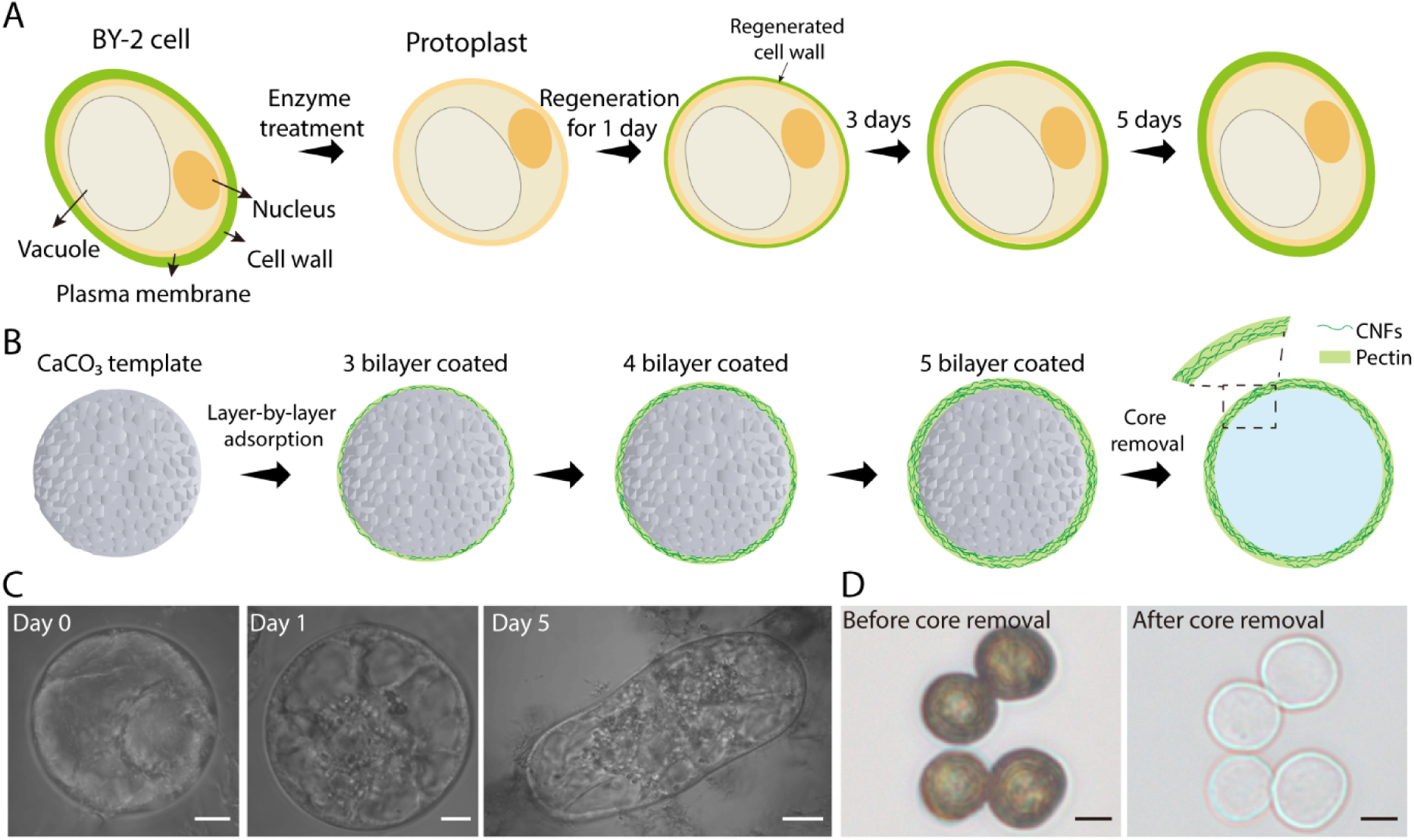
(A) Regeneration of cell wall in BY2 protoplasts and (B) bottom-up assembly of synthetic analogues by adsorption of pectin and CNFs layers on top of CaCO_3_ template particles. (C)The shapes of BY2 protoplasts after 0,1 and 5 days of cell wall regeneration. (D) The template of synthetic capsules was removed using acid and washed with MiliQ-water, yielding a core-shell structure with water-filled core and multilayered wall. Scale bars = 5 µm (C, D) or 10 µm (C, day 5).

## Notes

### Competing Interest Statement

The authors have declared no competing interest.

