## Supplementary Information for "Synthetic pectin-cellulose nanofiber capsule provides minimal model capturing mechanics of a regenerating plant cell wall"

**Table S1.** Quantification of monosaccharides for regenerated and synthetic walls
Values are presented as mean ± standard deviation.

| Monosaccharide | Synthetic wall (ng/mg DW) | Regenerated wall Day5 (ng/mg DW) |
|---|---|---|
| **Arabinose** | 33.49 ± 1.05 | 25.05 ± 11.02 |
| **Fucose** | n.a.* | n.a.* |
| **Galactose** | 382.78 ± 92.45 | 34.02 ± 18.89 |
| **Galacturonic acid** | 1899.07 ± 1063.90 | 5.61 ± 1.39 |
| **Glucuronic acid** | 8.01 ± 6.42 | 0.00 ± 0.00 |
| **Glucose** | 847.41 ± 286.04 | 40.77 ± 17.90 |
| **Mannose** | 83.76 ± 18.41 | 30.90 ± 9.55 |
| **Rhamnose** | 55.68 ± 18.66 | 1.97 ± 0.65 |
| **Xylose** | 94.77 ± 22.47 | 3.94 ± 0.16 |

*n.a. = not attained.

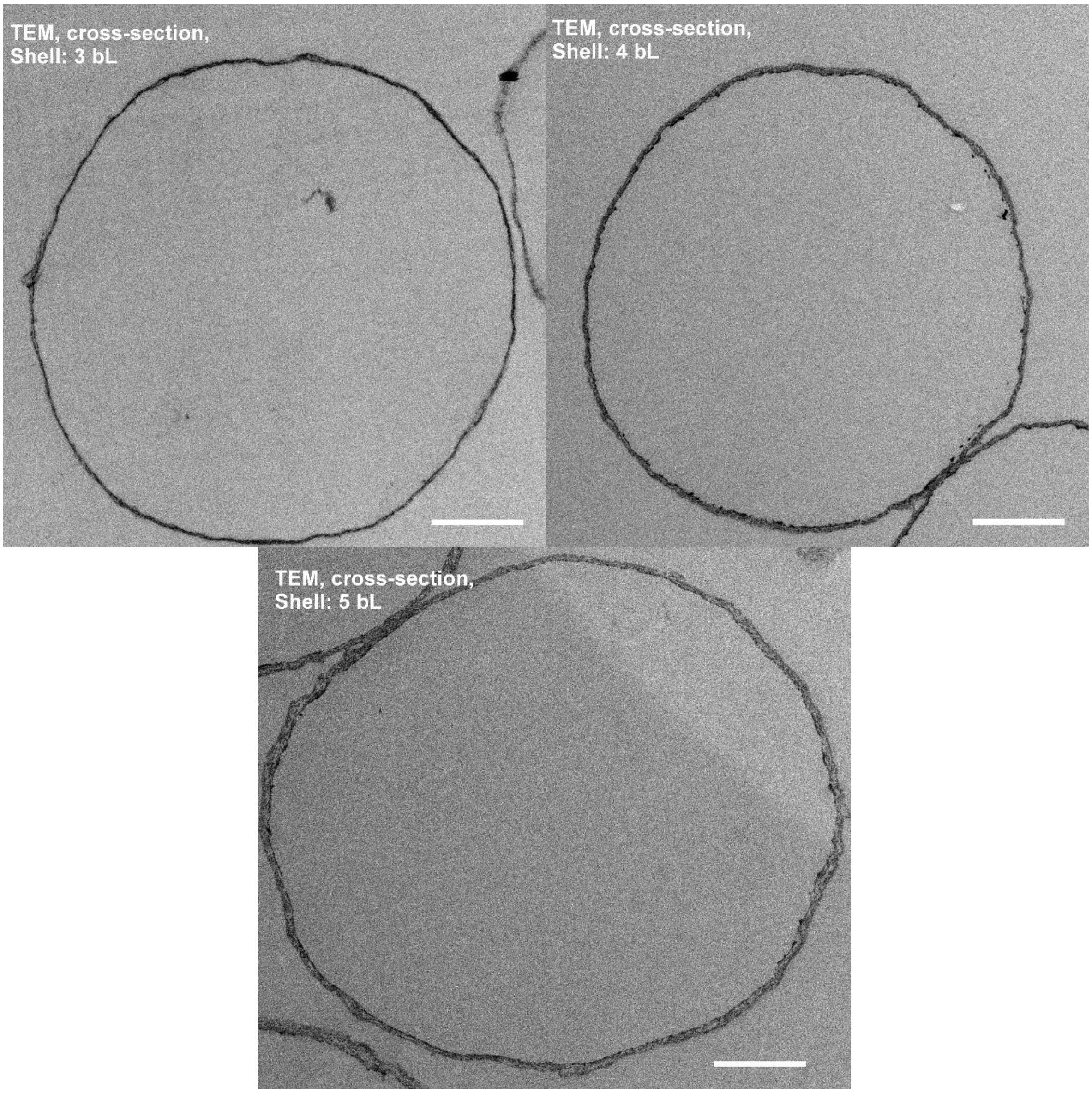


**Supplementary Fig 1**. TEM cross-sections of synthetic analogues with walls made from 3 bL, 4 bL and 5 bL. Scale bars=2 μm.

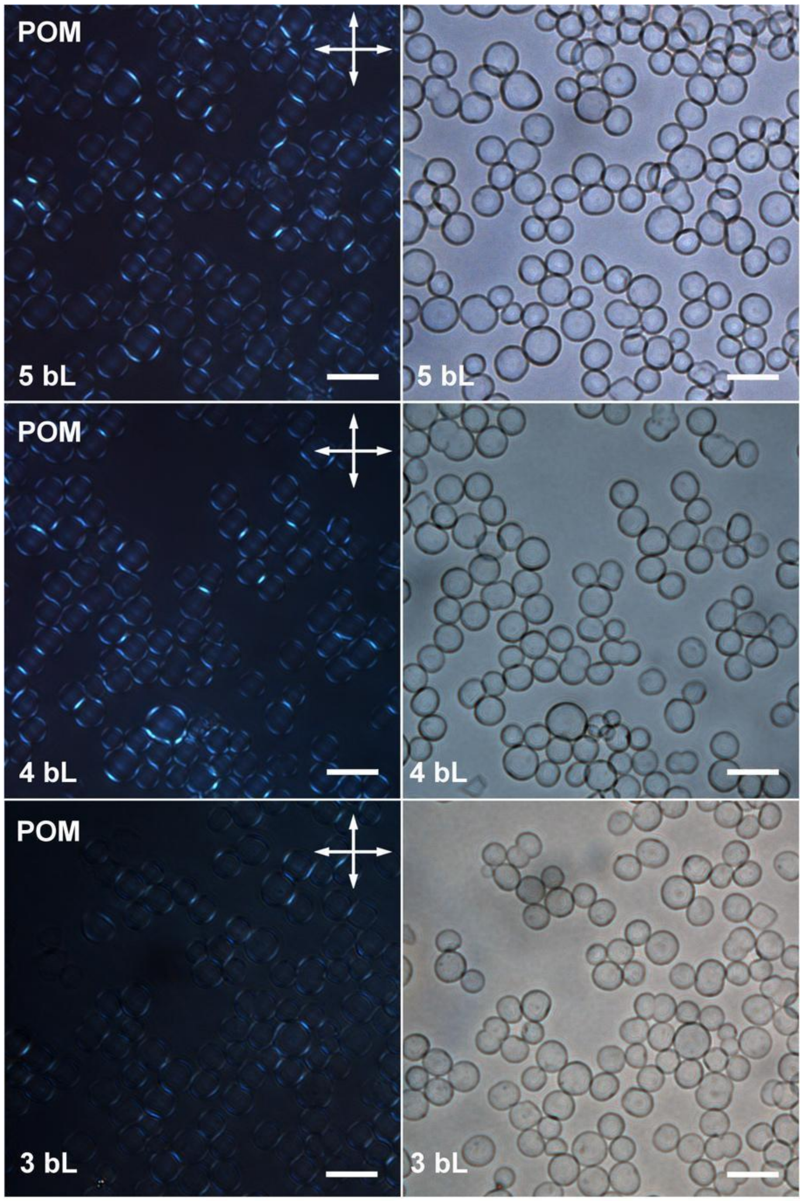


**Supplementary Fig 2**. Polarized optical microscopy (POM) and Bright field images of synthetic walls made from 3 bL, 4 bL and 5 bL of pectin and CNF. Scale bars=20 μm.

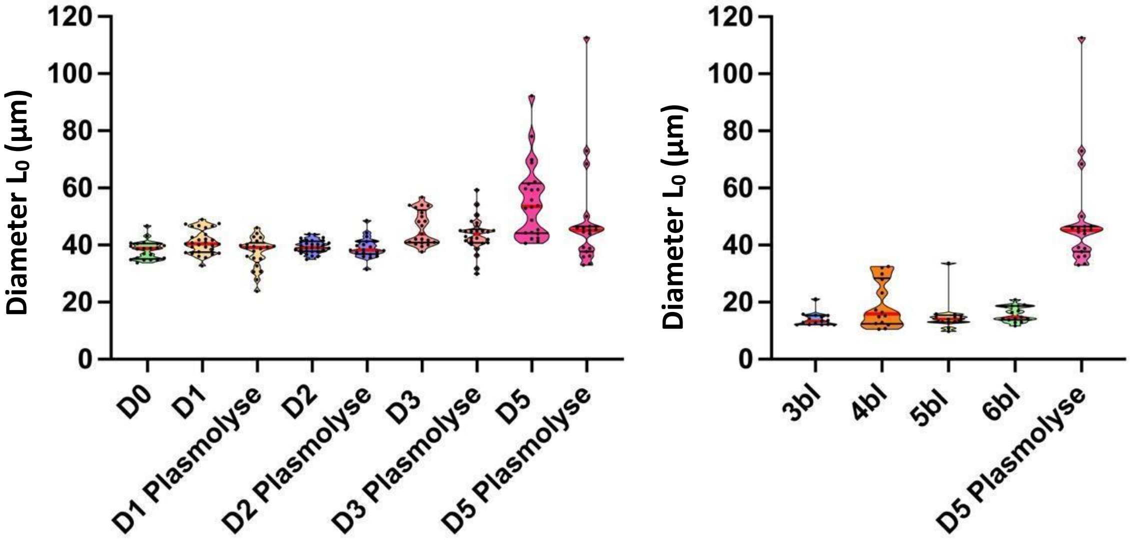


**Supplementary Fig 3.** The diameter of the cells and synthetic analogues, measured during microplate-based mechanical assays, was used to define the initial contact length ($L_0$) for the calculation of mechanical parameters. Notably, plant cells exhibit diameters at least three to four times greater than those of the synthetic analogues.
